# Co-release of histamine and GABA in prefrontal cortex excites fast-spiking interneurons and causes divisive gain change in pyramidal cells; an effect that is enhanced in older mice

**DOI:** 10.1101/2022.03.11.483936

**Authors:** Diana Lucaci, Xiao Yu, Paul Chadderton, William Wisden, Stephen G Brickley

## Abstract

We studied how co-release of histamine/GABA from axons originating from the hypothalamic tuberomammillary nucleus (TMN) and projecting to the prefrontal cortex (PFC) influences circuit processing. We opto-stimulated histamine/GABA co-release from genetically defined TMN axons that express the histidine decarboxylase gene (TMN_HDC_ axons). Whole-cell recordings were used to monitor excitability of visually identified PFC neurons in layer 2/3 of prelimbic (PL), anterior cingulate (AC) and infralimbic (IL) regions before and after opto-stimulated histamine/GABA release. We found that histamine-GABA co-release influences the PFC through actions on distinct neuronal types: histamine stimulates fast-spiking interneurons; and co-released GABA enhances tonic (extrasynaptic) inhibition on pyramidal cells (PyrNs). For fast spiking non-accommodating interneurons, opto-stimulation increased excitability, an effect blocked by histamine H1 and H2 receptor antagonists. The excitability of other interneuron types in the PFC was not altered. In contrast, the combined action of histamine and GABA co-release from TMN_HDC_ axons produced predominantly divisive gain changes in PyrNs, increasing their resting input conductance, and decreasing the slope of the input-output relationship. The direct inhibitory effect of TMN_HDC_ axon activation on PyrNs was not blocked by histamine receptor antagonists but was blocked by GABA_A_ receptor antagonists. Across the adult lifespan (from 3 months to over 2 years of age), stimulation of TMN_HDC_ axons in the PFC inhibited PyrN excitability significantly more in older mice. For individuals that maintain cognitive performance into later life, increases in TMN_HDC_ modulation of PyrNs could enhance information processing and be an adaptive mechanism to buttress cognition.

## Introduction

Histamine, a wake-specific neuromodulator, is produced in tuberomammillary nucleus (TMN) neurons of the posterior hypothalamus (Watanabe et al., 1983; Panula et al., 1984; Haas and Panula, 2003; Scammell et al., 2019; Yoshikawa et al., 2021). These neurons send axons throughout the brain (Takagi et al., 1986; Airaksinen and Panula, 1988; Haas and Panula, 2003; Arrigoni and Fuller, 2021; Yoshikawa et al., 2021), and are defined by expression of the histidine decarboxylase (*hdc*) gene, which encodes the enzyme that synthesizes histamine (Joseph et al., 1990). Histamine is released from axonal varicosities (Takagi et al., 1986), and activates excitatory H1, H2 and inhibitory H3 metabotropic receptors (Haas and Panula, 2016). As histamine acts by volume/paracrine transmission, it influences multiple elements of the circuitry, fine-tuning both inhibitory and excitatory transmission (Ellender et al., 2011; Bolam and Ellender, 2016). Increased firing of neurons in the TMN is associated with the wake state of an animal and antihistamines that cross the blood brain barrier cause drowsiness. Therefore, it is expected that histamine release within the neocortex will result in excitation of the cortical circuitry.

Dual transmitter release is a common phenomenon, e.g. GABA-acetylcholine, GABA-dopamine, and glutamate-orexin co-release have all been documented (Ma et al., 2018). Similarly, histamine neurons contain GABA as well as the enzyme glutamate decarboxylase GAD1 (GAD67) needed for GABA synthesis (Takeda et al., 1984; Senba et al., 1985; Airaksinen et al., 1992; Trottier et al., 2002). For GABA transport into synaptic vesicles, the VGAT protein (encoded by the *vgat/slc32a1* gene) is usually needed (McIntire et al., 1997). Co-localization of *vgat* and *hdc* transcripts (Abdurakhmanova et al., 2020), as well as evidence from *hdc-Cre* x *vgat* reporter mouse crosses (Yu et al., 2015), and visualization of GABA in vesicles of histaminergic cells *in vitro* (Kukko-Lukjanov and Panula, 2003), suggested GABA release from vesicles in histamine neurons (Ma et al., 2018). Indeed, we found that histaminergic axons broadcast both GABA and histamine (Yu et al., 2015).

As well as causing fast (millisecond) synaptic inhibition through GABA_A_ receptors, GABA can activate to extrasynaptic ionotropic GABA_A_ receptors to produce a tonic shunting inhibition (Brickley et al., 1996; Brickley et al., 2001; Wall, 2002; Brickley and Mody, 2012; Lee and Maguire, 2014; Brickley et al., 2018). Knockdown of *vgat* in *hdc* cells enhanced activity of mice and reduced their sleep. In the visual cortex, light-evoked GABA release increased tonic inhibition onto pyramidal cells, which was diminished when *vgat* gene expression was knocked down (Yu et al., 2015). Therefore, our work has challenged the view that histaminergic neurons’ influence on cortical circuitry will be purely excitatory.

V*gat* expression in histaminergic cells has become controversial, however. Combined immunocytochemistry and *in situ* hybridization for HDC protein and *vgat* mRNA revealed little co-expression (Venner et al., 2019) and single-cell RNA seq studies of mouse TMN identified a subset (E5) of *hdc* cells with moderate expression of *vgat* (Mickelsen et al., 2020). Nevertheless, this subset of cells due to their long-range projections, could still have widespread influence. Therefore, in this study we have extended our analysis to the prefrontal cortex (PFC) and broadened our investigation to assay excitability changes in pyramidal cells and interneurons. We have used whole-cell recording techniques combined with optogenetics to describe how the gain of the input-output relationship is altered following GABA/histamine release from hdc expressing TMN axons (TMN*_HDC_* axons). Importantly, we have now recorded over a much broader age range than in our previous study (Yu et al., 2015) to ask whether the changes in excitability associated with GABA/histamine release are uniform across the adult lifespan. For the first time, we show that GABA release is elicited from genetically defined histaminergic axons in the PFC of young and old mice and that the GABA/histamine cotransmission works together to enhance pyramidal cell information transfer, especially in older animals.

## Materials and Methods

### Mice

All experiments were performed in accordance with the United Kingdom Home Office Animal Procedures Act (1986) and were approved by the Imperial College Animal Welfare Ethical Review Committee. The strains of mice used were HDC-Cre (*Hdc^tm1.1(icre)Wwis/J^*, JAX stock 021198) (Zecharia et al., 2012) and Pv-Cre (*Pvalb^tm1(cre)Arbr/J^*, JAX stock 008069), kindly provided by S. Arber (Hippenmeyer et al., 2005). The *HDC-Cre* mouse line contains an *ires-Cre* cassette knocked-in to exon 12 of the *hdc* gene. The insertion is downstream of the stop codon for the *hdc* reading frame, and therefore the knock-in allele makes HDC and Cre proteins (Zecharia et al., 2012).

### AAV vectors and surgery

AAVs (capsid serotype 1/2) were packaged in-house, and contained either an *AAV-EF1a-flex-ChR-ChR2H134R-EYFP* transgene (a gift from Karl Deisseroth, Addgene plasmid 20298) or an *AAV-CBA-flex-GFP* transgene (a gift from Edward Boyden, Addgene plasmid 28304). To deliver the *AAVs* into the brain, bilateral stereotaxic injections were performed on *HDC-Cre* or *Parv-Cre* mice using an Angle Two apparatus (Leica) linked to a digital brain atlas (Leica Biosystems Richmond, Inc.). Before injections, 1 μl AAV virus was mixed with 1 μl 20% mannitol (MERCK K93152782 111). The virus and mannitol mixture were injected into a pulled-glass pipette (Warner Instruments; OD = 1.00 mm, ID = 0.78 mm, length = 7.5 cm). Virus was injected at a speed of 25 nl/min. Virus (1 μl) was bilaterally injected into the brain, 0.5 μl for each side. For the TMN, the injection coordinates were: ML (−0.92 mm), AP (−2.70 mm), DV (−5.34 mm); ML (0.92 mm), AP (−2.70 mm), and DV (−5.34 mm) (Yu et al., 2015). For the frontal cortex the coordinates were: ML (0.33 mm), AP (2.10 mm), DV (−2.13 mm), and 0.3 μl of virus was injected.

After each injection, pipettes were left undisturbed for 10 min to allow the viral solution to be absorbed by the tissue; thereafter, pipettes were removed slowly. After injection, craniotomies were sealed with Kwik-Cast (World Precision Instruments, USA), and the scalp was sutured with glue (Histoacryl, Braun, USA) or nonabsorbable nylon sutures (Ethilon, Ethicon, USA). Immediately after suturing, anesthesia was antagonized via injection of a naloxone, flumazenil, and atipamezole mixture (1.2, 0.5, and 2.5 mg/kg, i.p.). Postoperative analgesia was provided via eutectic mixture of local anesthetics (EMLA; 2.5% lidocaine and prilocaine) application to the sutured skin. Mice were placed in a heated recovery box under observation until they fully woke up from anesthesia; thereafter, animals were returned to their home cage. Four to five weeks after the AAV injection, we prepared acute slice preparations containing the TMN area. If there was visible primary fluorescence in this area we also went on to slice the PFC. Initially, we also tested for the presence of ChR2 by patching the cells fluorescent TMN cells, and in current clamp mode and eliciting an inward current response with blue light stimulation of ChR2 to confirm the presence of functional ChR2. Slices from TMN injections that gave no visible primary expression were discarded or used as negative controls.

### PFC Slice Electrophysiology

Acute brain slices were prepared following cervical dislocation. Each mouse brain was rapidly removed from the cranium and immediately immersed in ice cold, oxygenated slicing solution. The slicing solution contained (in mM: NMDG 92, KCl 2.5, NaH2PO4 1.25, 30 mM NaHCO3, HEPES 20, glucose 25, thiourea 2, Na-ascorbate 5, Na-pyruvate 3, CaCl2·4H2O 0.5 and MgSO4·7H2O 10) pH 7.3-7.4 when bubbled with 95%O2/5%CO2 and titrated with concentrated hydrochloric acid. Acute slice preparations were cut using a vibratome (Campden instruments) at a thickness of 250 μm, after which they were immediately transferred to a holding chamber containing slicing ACSF at 33-34°C continuously bubbled with 95%O2/5%CO2, Slices were kept in this condition for a 10-15 minutes, after which they were transferred into a holding chamber at room temperature containing recording ACSF (in mM: NaCl 125, KCl 2.5, CaCl2 2, MgCl2 1, NaH2PO4 1.25, NaHCO3 26, glucose 11, pH 7.4 when bubbled with 95%O2/5%CO2) that was continuously bubbled with 95%O2/5%CO2.

Slices were then visualized using a fixed-stage upright microscope (BX51W1, Olympus and Scientifica Slice scope) fitted with a high numerical aperture water-immersion objective and an infra-red sensitive digital camera. Patch pipettes were made from thick-walled borosilicate glass capillaries (0.86 mm internal diameter, 1.5 mm outer diameter, Harvard Apparatus) using a two-step vertical puller (Narishige, PC-10). Pipette resistances were typically 3-4 MΩ when back filled with internal solution. For voltage-clamp experiments, the internal solution contained (in mM): CsCl 140, NaCl 4, CaCl2 0.5, HEPES 10, EGTA 5, Mg-ATP 2; the pH was adjusted to 7.3 with CsOH. For current-clamp experiments the internal solution contained (in mM): 145 K-gluconate; 4 NaCl; 0.5 CaCl2; 10 HEPES; 5 EGTA; 4 Mg-ATP; 0.3 Na-GTP (adjusted to pH 7.3 with KOH). The amplifier head stage was connected to an Axopatch 700B amplifier (Molecular Devices; Foster City, CA). The amplifier current output was filtered at 10 kHz (–3 dB, 8-pole low-pass Bessel) and digitized at 20 kHz using a National Instruments digitization board (NI-DAQmx, PCI-6052E; National Instruments, Austin, Texas). Data acquisition was performed using CED Power3 1401 processor and CED Signal (Version 6) software.

### Optogenetics

A 470 nm collimated LED (M470L3-C1, Thorlabs) was used to illuminate the slice through the objective lens. The LED was driven by an LED-driver (LEDD1B, Thorlabs) which was controlled by a National Instruments digitization board (NI-DAQmx, PCI-6052E; National Instruments, Austin, Texas). The output of the LED was measured with a power meter (PM100D, Thorlabs) positioned below the objective lens and the power output and membrane conductance or voltage changes were aligned for analysis. The optical power emitted through our 63X water immersion lens increased linearly to a maximum power of 70 mW/mm^2^ at our chosen light stimuli, giving rise to a transient response that peaked at 40 mW/mm^2^, with a 10%–90% rise time of 0.73 ms and a decay constant of 9.65 ms.

### Data analysis

For all recorded cells, total membrane capacitance (**C_m_**) was calculated in voltage-clamp configuration from Cm = Q/ΔV, where Q was the charge transfer during a hyperpolarizing 10 mV step of the command voltage (ΔV). The total membrane conductance (**G_m_**) was calculated from G_m_ = I_ss_/ΔV, where I_ss_ was the average steady-state current during the ΔV. The electrode-to-cell series resistance (**Rs**) was calculated from the relationship Rs = ΔV/I_p_, where I_p_ was the peak of the capacitive current transient and recordings were excluded if Rs increased by >30%.

### Experimental Design and Statistical Analysis

To test for statistical significance between single-cell properties across our broad range of ages, we first applied the Mann-Whitney non-parametric test. We then applied the Benjamini-Hochberg (B-H) correction at p= 0.05 and 0.01 values to control for multiple comparisons. As we were looking for changes in many biophysical properties we chose the B-H correction procedure as it is less sensitive than the Bonferroni procedure to decisions about the identity of a “family” of tests. Briefly, the B-H procedure ranks individual p-values and then compares individual p-values to their B-H critical value, (*i/m*)\**Q*, where *i* is the rank, *m* is the total number of tests, and *Q* is the false discovery rate. The largest p-value that has *p* < (*i/m*)\**Q* is considered significant.

## Results

Four to five weeks prior to whole-cell recording experiments, an AAV containing a flex-ChR2-EYFP transgene was delivered into the TMN of HDC-Cre mice (**Figure 1A**). The expression of ChR2-EYFP was verified in the TMN (**Figure 1B**) and, like our previous reports, we found that ChR2-EYFP expression was restricted to a subset of hdc expressing neurons of the TMN (**Figure C**). Whole-cell recordings were then made from fast-spiking interneurons (FS-Ins) and pyramidal neurons (PyrNs) within layer 2/3 of the prelimbic (PL), anterior cingulate (AC) and infralimbic (IL) regions of the PFC (**Figure 1D**). Following these recordings, the presence of fluorescently labelled TMN_HDC_ axons was observed in the PFC (**Figure 1E**). 3D reconstructions of biocytin filled cells in the PFC, demonstrated how these fluorescently labelled TMN_HDC_ axons rarely make close appositions (< 1μm) with PyrN dendrites (**Figure 1F**), as expected from previous studies (Takagi et al., 1986). Pyramidal neurons (PyrNs), fastspiking accommodating (FS-INa) and fast-spiking non-accommodating interneurons (FS-INna) in layer 2/3 were selected according to their location, soma shape, and electrophysiological features. Most strikingly, PyrNs exhibited evoked AP firing rates of below 20 s^-1^, with an average maximum AP rate of 11.4 ± 0.5 s^-1^ (n = 19), whereas the evoked firing rates of FS-INs was between 20 s^-1^ to 90 s^-1^ with an average maximum AP rate of 46.6 ± 3.8 s^-1^ (n = 19) (**Figure 1G**). To further confirm the identity of FS-INs, we patched GFP-fluorescent cells from Parv-Cre mice following AAV-flex-GFP injection into the PFC (n = 4). As expected, AP characteristics for the parvalbumin-expressing cells were like those observed in the FS-IN group. A further breakdown of the FS-IN group was based on differences in the inter spike intervals (ISIs) of FS-INs (**see Figure 1H**). Cells with stable ISIs were classified as nonaccommodating FS interneurons (FS-IN_na_) whereas fast-spiking cells with an unstable ISI were classified as accommodating FS interneurons (FS-INa). This variability score was close to zero for FS-IN_na_ cells (Figure 1I) and there was no overlap with the ISI variability in the FS-INa population (FS-IN_a_= 11, FS-IN_na_= 15, p < 1.3E-7 by a Mann-Whitney U-test). The regular fast spiking interneuron population (FS-IN_na_) could contain basket, chandelier and neurogliaform cell types that all provide axo-somatic feed-forward and feed-back inhibition onto PyrNs (Feldmeyer et al., 2018). Based upon firing properties alone, the irregular spiking population (FS-IN_a_) are likely to represent the somatostatin positive Martinotti and non-Martinotti cells that target inhibition to the basal and apical dendrites of PyrNs (Feldmeyer et al., 2018). However, additional morphological and immunohistochemical analysis would be required to confirm this classification (Krimer et al., 2005).

**Figure 1.**
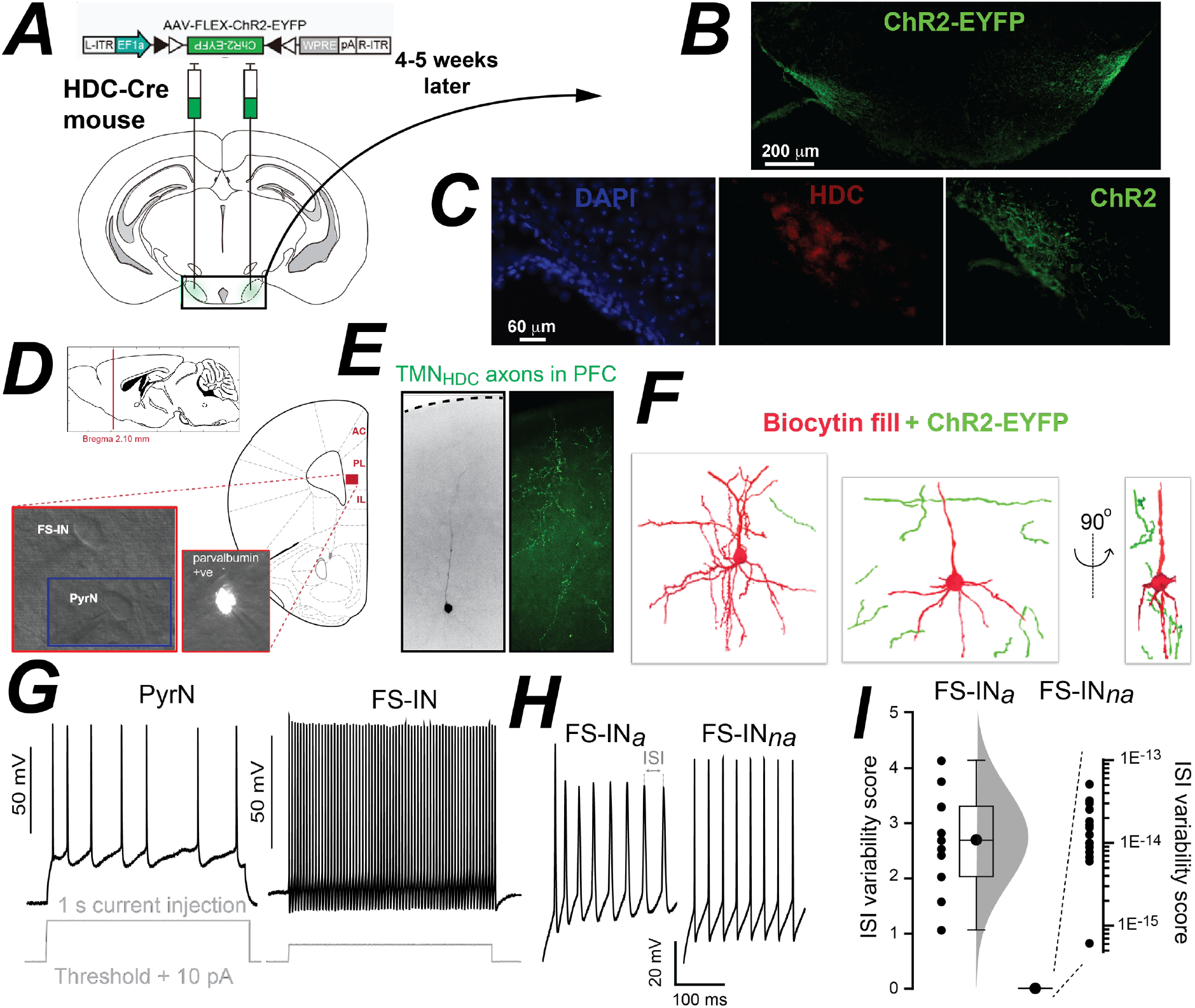
Targeting of ChR2-EYFP expression to TMN axons and Identification of neuronal types in the PFC. ***A,*** Depiction of the construct used to target ChR2-EYFP expression to neurons of the TMN in the HDC-Cre mouse strain. The location of bilateral AAV injections is shown superimposed on an image taken from the Allan Brain Atlas. **B,** Low magnification epifluorescent image of a coronal brain section taken 4-5 weeks after AAV injection demonstrating EYFP expression in the TMN. ***C,*** Higher magnification image of all cell bodies present in the TMN on one side of the brain using a nuclear DAPI stain (blue) and a comparison with histamine expressing neuron identified using the HDC antibody (red) and the ChR2 expressing neurons labelled as in ***B. D,*** Illustration of the PFC region where, 4-5 weeks later, blue light was delivered through a high numerical aperture imaging objective to stimulate ChR2. Included in this panel is a sagittal location of the mouse prefrontal cortex (PFC) along with a coronal section showing the three areas of the PFC where recordings were made: anterior cingulate (AC), prelimbic cortex (PL) and infralimbic cortex (IL). The illustrations were adapted from the Allen Brain Mouse Atlas. Bright field images were taken during whole cell recording and the fluorescent cell was recorded from the Parv-Cre mouse following flexed-GFP delivery into the PFC. ***E,*** Microscope images taken from the PFC region where recordings were made, and blue light was delivered. The lefthand bright field image demonstrates the location of the biocytin filled PFC neuron and the epifluorescent image on the right illustrates the presence of ChR2-EYFP labelled axon terminals in this region of the PFC. ***F,*** PFC neurons were reconstructed following confocal microscopy to reveal the proximity of the TMN axons relative to the recorded neuron. As expected, the ChR2-EYFP labelled axon terminals were never observed close enough (<1μm) to indicate the presence of conventional synaptic connections. ***G,*** Examples of voltage recordings made in whole-cell current-clamp configuration during steady-state 1 second current injections sufficient to elicit APs. A representative voltage trace is shown from a pyramidal neuron (PyrN) and a fast-spiking interneuron (FS-IN). ***H***, Comparison of accommodating (FS-IN_*a*_) and non-accommodating (FS-IN_*na*_) FS-IN firing. ***I,*** A plot of the ISI changes for the two recordings shown in ***H***. Comparison of inter-spike interval (ISI) analysis standardised to zero mean and 1 standard deviation. The sum of the standardised ISI is referred to as the ‘ISI variability score’. This score is extremely close to zero for the FS-IN_na_ group, hence the logarithmic scale. No overlap between the groups is observed. Difference in ISI score is significant to p < 0.01 using Mann Whitney and Benjamini-Hochberg correction.

### Optogenetic stimulation of TMN_HDC_ axons alters excitability in the PFC

During whole-cell recording, TMN_HDC_ fibres in the PFC slices were activated with 1 ms duration blue light flashes delivered at a rate of 5 s-1 every 0.4 seconds for a total stimulation period of 4 minutes (Yu et al., 2015). In the example shown in **Figure 2A**, simultaneous whole-cell recordings were made from a FS-IN_*na*_ and a PyrN. In PyrNs, a reduction in AP firing rates following TMN_HDC_ axon stimulation was obvious from the input-output (I-O) relationships constructed from average AP rates. These I-O relationships were well described by a sigmoidal function. Results from the fits obtained from 19 PyrNs were pooled, and the average I-O relationship before and after optogenetic stimulation of TMN_HDC_ axons was constructed (**Figure 2B**). This analysis demonstrated that the maximum AP firing rate was reduced from 12.7 ± 1.6 s^-1^ to 3.9 ± 5.3 s^-1^ following stimulation of TMN_HDC_ axons (Wilcoxon signed rank test, p=0.0002). The slope of the average I-O relationship significantly reduced from 7.6 ± 0.9 to 2.4 ± 0.7 (Wilcoxon signed rank test, p=0.0004) with only a small reduction in the current required to reach 50% of the maximum AP rate (50.4 ± 36.2 pA before stimulation and 49.9 ± 18.9 pA after stimulation) that was not significant (Wilcoxon signed rank test, p=0.7). Transforming the average, I-O relationship obtained under control conditions with a purely divisive function indicates that an additive gain change mechanism may make a minor contribution to changes in PyrN excitability. Specifically, there was a small left-ward shift in the current required to reach AP threshold that was not predicted from a purely divisive model (**Figure 2B**). However, the most striking aspect of the changes in the I-O relationships constructed from PyrN recordings were the divisive effects observed following TMN_HDC_ axon stimulation.

**Figure 2.**
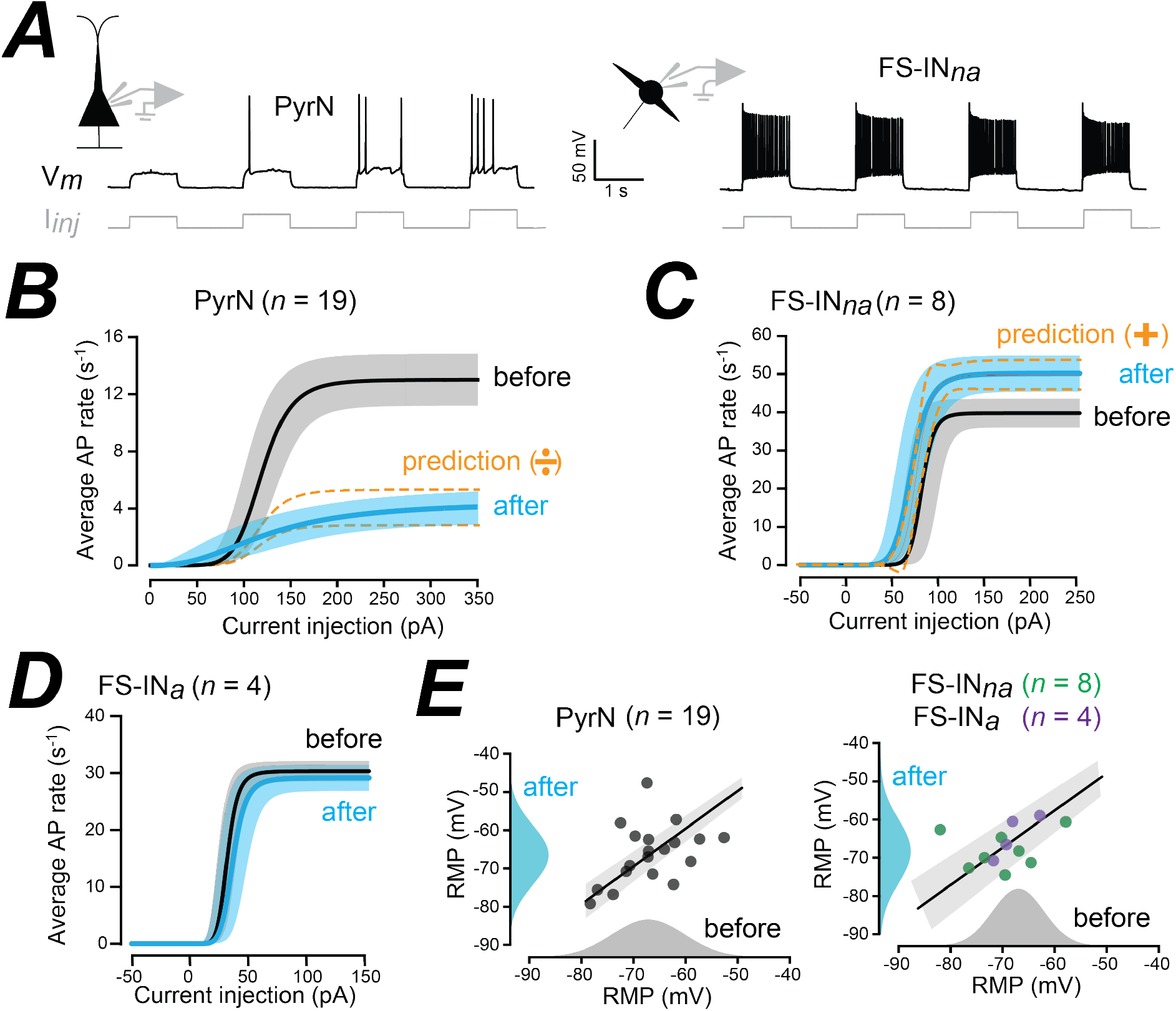
Optogenetic release of GABA/Histamine from axons within the adult PFC. ***A,*** Voltage traces obtained during simultaneous whole-cell recording from a PyrN and a FS-IN_*na*_ used to monitor changes in AP firing. ***B,*** Comparison of the average fits obtained from the input-output relationships constructed from the PyrN population (n=19) before and after optogenetic stimulation of the TMN_HDC_ axon terminals in the PFC. The solid black line is the average fit with the grey shaded area showing the 95% confidence limits for the fits obtained before stimulation. The blue solid line is the average fit and the shaded blue region is the 95% confidence limits for fits obtained after optogenetic stimulation. The dashed lines demonstrate the predictions when assuming a purely divisive gain change mechanism in PyrNs. ***C,*** Comparison of the average fits obtained from the input-output relationships constructed from the FS-IN*_na_* population (n=8) before and after optogenetic stimulation of the TMN_HDC_ axon terminals in the PFC. In contrast to the data from PyrNs, an additive gain-change mechanism was solely responsible for the increased firing observed in the FS-IN_na_ population. ***D***, Comparison of the average fits obtained from the input-output relationships constructed from the FS-IN_*a*_ population (n=4) before and after optogenetic stimulation of the TMN_HDC_ axon terminals in the PFC. Note the lack of any change in the FS-IN_a_ population. ***E***, Scatter plots for the RMP recorded before and after optogenetic stimulation in PyrNs and FS-INs with the results of linear region analysis. The all-point histograms constructed before and after optogenetic stimulation further demonstrates how the RMP does not alter. The straight lines are the results of linear regression analysis with the 95% confidence limits shown in grey shaded areas.

The average firing rates of the FS-IN*na* population (*n*=8) significantly increased following optogenetic stimulation of TMN_HDC_ axons (**Figure 2C**). The change in the average I-O relationship of the FS-IN*na* population was consistent with an additive gain change mechanism (**Figure 2C**). The maximum AP firing rate increased from 39.8 ± 3.8 s^-1^ to 44.8 ± 3.8 s-1 (Wilcoxon signed rank test, p=0.008) with 50% excitation point shifting from 82.4 ± 17.8 pA to 70.0 ± 16.2 pA (Wilcoxon singed rank test, p=0.5). However, there was also a small but significant reduction in the slope of the input-output relationship from 0.3 ± 0.2 to 0.2 ± 0.1 (Wilcoxon signed rank test, p=0.008). Therefore, like the situation for PyrNs, analysis of I-O relationships suggests that two types of gain mechanism could operate within the FS-IN_na_ population. However, transforming fits from the FS-IN_na_ population solely with an additive function predicted well the changes observed in the I-O relationship.

In contrast to the clear changes observed for PyrNs and FS-IN_na_s, the excitability of the FS-IN*_a_* population was unchanged following optogenetic stimulation of TMN_HDC_ axons in the PFC (*n*=4), such that the maximum AP rate was 30.4 ± 1.8 s-1 before stimulation and 29.2 ± 2.2 s-1 after. There was also no significant change in the slope of the I-O relationship (6.8 ± 1.4 versus 6.4 ± 1.4) or the current required to reach 50% of the maximum AP rate (28.2 ± 7.1 pA versus 32.9 ± 11.1 pA) in the FS-IN*_a_* population (**Figure 2D**).

Consistent with an additive gain change mechanism neither the resting membrane potential or the input conductance of the FS-IN_na_ population was altered following stimulation of TMN_HDC_ axons (**see figure 2G**). The resting membrane potential of the PyrN population was also not altered during the tonic shunting inhibition associated with this divisive gain change. However, consistent with a divisive gain change mechanism, there was a significant 20% increase in membrane conductance in PyrNs following optogenetic stimulation that was consistent with a shunting inhibition (see Figure 5E) (n_pyr_ = 22, p < 0.02 using paired sample Wilcoxon Signed rank test). The divisive gain change observed in PyrNs is consistent with GABA binding to extrasynaptic GABA_A_ receptors, whereas the additive gain change associated with the FS-IN*_na_* population is more likely to be due to modulation of excitability following the action of histamine on G-protein coupled receptors. To test this hypothesis, we next undertook some simple pharmacological experiments, during which the time course of the gain change was also examined.

### The PyrN divisive gain change involves GABA_A_ receptors

To compare the time course associated with changes in PyrN excitability the average AP rate was calculated during the entire depolarizing current injection protocol (**Figure 3A**). The subthreshold membrane voltage appeared more linear following TMN_HDC_ axon activation and in 10 out of 22 PyrNs tested, no APs could be elicited following optogenetic stimulation. However, with larger depolarising currents, AP firing was possible in these PyrNs (data not shown). Simple Boltzmann functions fitted to the time course plots demonstrated that the reduction in PyrN firing was apparent at 2-3 minutes following TMN_HDC_ axon activation (**Figure 3B**). Analysis of all fits demonstrated that the average AP rate reduced significantly from 10.7 ± 0.8 s^-1^ to 4.5 ± 0.9 s^-1^ following optogenetic stimulation (n=22) and the average change in the AP rate for this group was −65.7% ± 8.4%. Several negative control experiments were conducted with acute PFC slices from *HDC-Cre* mice (*n* = 8) which had not received *AAV-flex-ChR2* injections into the TMN. In these experiments, the optogenetic protocol resulted in a −15.9% ± 7.0% change in AP firing rates in PyrNs that was not significant (n = 8, p < 0.06, Wilcoxon Signed Ranks Test). There was also no change in resting membrane potential or membrane conductance in these PyrNs. As a further control, the addition of the GABA_A_ receptor antagonist SR95531 blocked the actions of TMN_HDC_ axon stimulation, resulting in only a small reduction in AP firing rates in PyrNs of −7.7% ± 10% following optogenetic stimulation that was also not significant (n_H1,H2_ = 5, n_GABA-A_ = 5, p < 0.029 by Mann Whitney).

**Figure 3.**
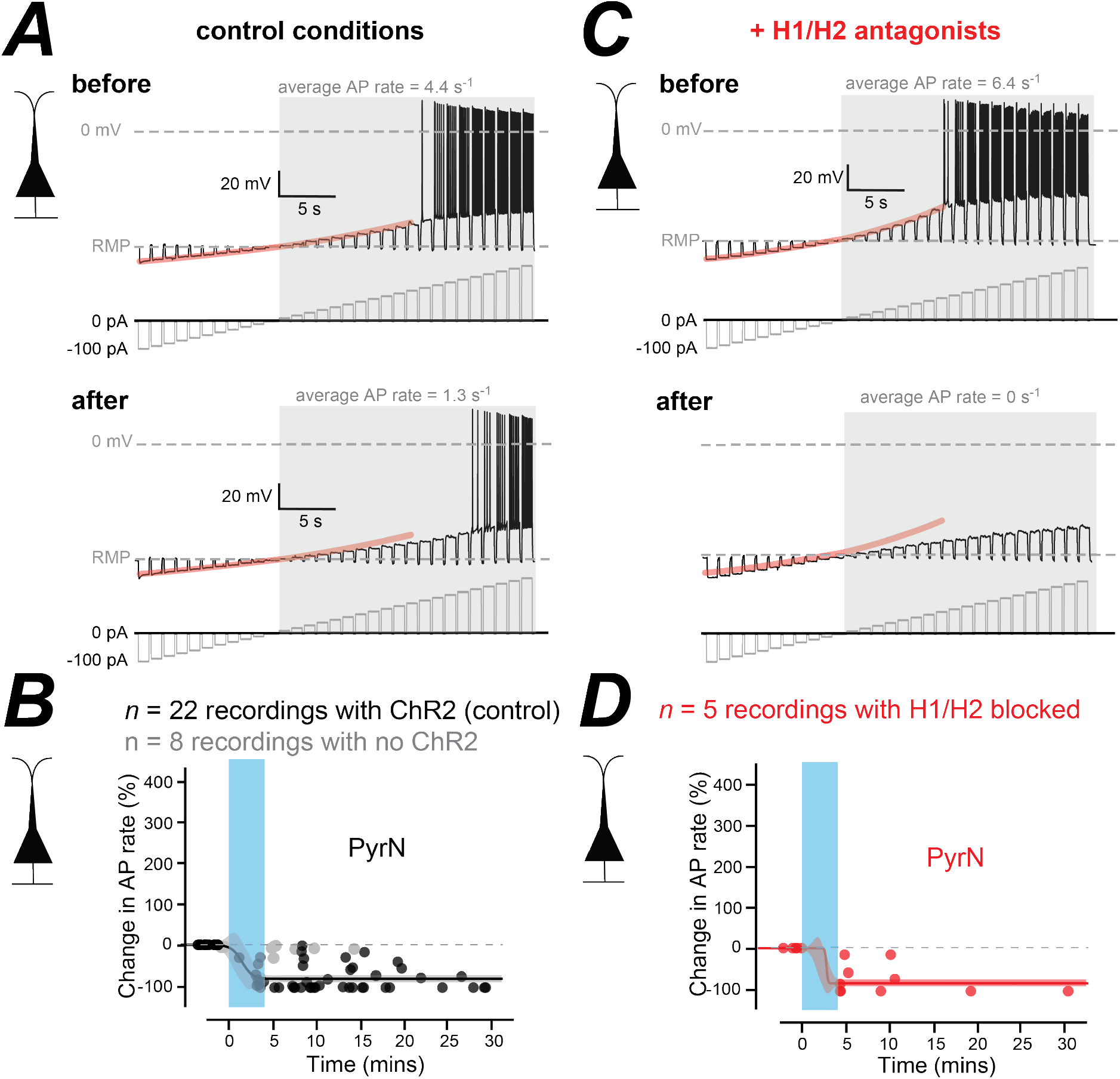
The effect of blocking H1/H2 receptors on changes in the excitability of the PyrNs in the PFC. ***A***, Voltage traces recorded from a PyrN, before and after optogenetic stimulation of TMN_HDC_ axons. The current injection protocols are shown below each voltage trace in grey. The average AP rate was calculated during the shaded epoch of the recording following threshold crossing at 0 mV. The subthreshold membrane voltage relationship obtained before blue light stimulation is highlighted in orange and superimposed onto the voltage response recorded after blue light stimulation for comparison. In this case the subthreshold I-V relationship becomes more linear following light stimulation. ***B***, Plots illustrating the change in average AP rate in the presence (black symbols) and absence (grey symbols) of ChR2 expression in TMN_HDC_ axons. The time course of these changes in average AP rate can be described using a Boltzmann function to quantify the rate as well as the magnitude of the change in excitability. The shaded area indicates the 95% confidence limits for these fits. ***C*** & ***D,*** same conventions as ***A*** & ***B*** but the data was obtained from PyrNs in the presence of H1/H2 antagonists. Note in ***C*** how the subthreshold response of PyrNs becomes more linear following optogenetic stimulation of TMN_HDC_ axons and how in ***D*** the magnitude of the reduction in AP rate is like control conditions.

Pharmacological experiments were undertaken to directly address the involvement of H1/H2 receptors in the gain change. The optogenetic experiments described above were repeated in the presence of H1 (pyrilamine) and H2 (ranitidine) receptor antagonists. Immediately after the TMN_HDC_ axon stimulation protocol, a larger depolarising current was required to elicit APs in PyrNs (**Figure 3C**) and, on average, the AP firing rate in PyrNs decreased by −70.0% ± 15.8% in the presence of H1/H2 antagonists (n=5, p < 0.032, Wilcoxon Signed Ranks Test). In the example shown in **Figure 3C**, the average AP rate was 6.4 s-1 prior to blue light stimulation but was zero following stimulation. Therefore, in this PyrN the cell excitability was considered to have been reduced by −100% (**see Figure 3D**). Clear changes in the subthreshold voltage behaviour of this PyrN was still observed following TMN_HDC_ axon stimulation. The net inhibition of PyrNs observed following TMN_HDC_ axon stimulation in the PFC was −65.7% ± 8.4% (n=22) in control conditions compared to −70.0% ± 15.8% (n=5) in the presence of H1/H2 antagonists. A similar fast time course for PyrN AP firing rate changes were also observed in the presence of the H1/H2 blockers.

### The FS-IN_na_ additive gain change involves histamine receptors

Once again, the excitability of the FS-IN_a_ population was unchanged following optogenetic stimulation of TMN_HDC_ axons in the PFC but, the FS-IN_na_ population became more excitable (**Figure 4A**). In this example the average AP rate increased from 10.6 s^-1^ to 171.7 s^-1^ and this additive gain change was not associated with any change in the resting input conductance as evidenced by the linear subthreshold IV relationship before and after stimulation of TMN_HDC_ axons. The time course plots demonstrated that the enhancement of FS-IN*_na_* firing was noticeably delayed relative to the end of the blue light stimulation (**Figure 4B**) with the increase in AP rate occurring 10 minutes after the end of blue light stimulation at a much slower rate than that apparent for the reduction in PyrN excitability. This slow time course of the increase in AP firing could reflect a more gradual increase in histamine concentrations in the extracellular space or a mechanism involving modulation by G-protein-coupled receptors such as the H1/H2 receptors, whereas the rapid time course of the reduction in AP firing in PyrNs is consistent with the rapid binding of GABA to high-affinity ligand-gated ion channels. To examine the contribution of H1/H2 receptors in the additive gain change the optogenetic experiments described above were repeated in the presence of H1 (pyrilamine) and H2 (ranitidine) receptor antagonists. With histamine receptors blocked there was no change in the average AP rate after the TMN_HDC_ axon stimulation s (**Figure 4C**) and, on average, the AP firing rate in the FS-IN_na_ population was not increased in the presence of H1/H2 antagonists (**Figure 4D**).

**Figure 4.**
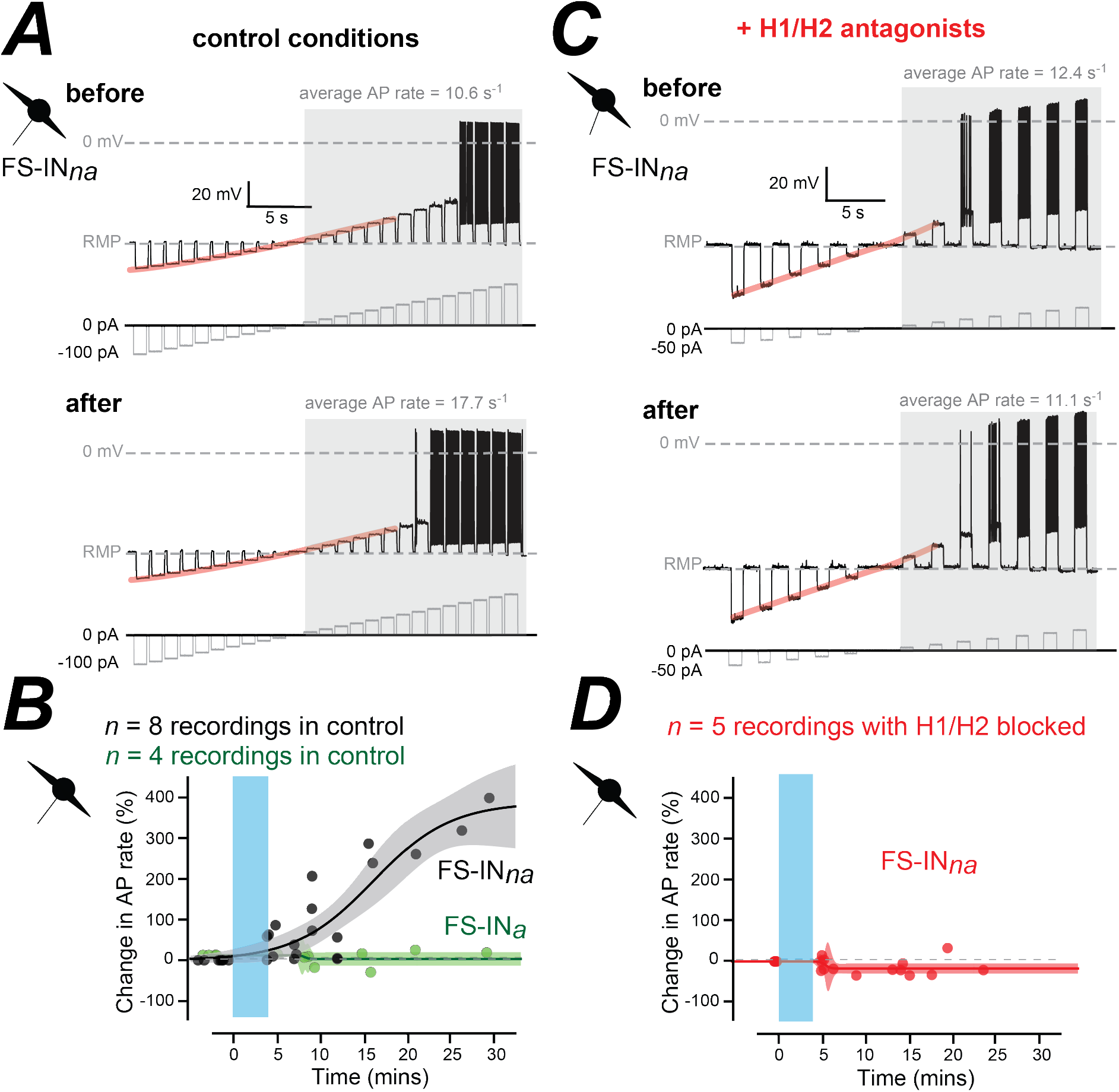
The effect of blocking H1/H2 receptors on changes in the excitability of the FS-IN_na_ population in the PFC. ***A***, Voltage traces recorded from a FS-IN_*na*_, before and after optogenetic stimulation of TMN_HDC_ axons. The current injection protocols are shown below each voltage trace in grey. The average AP rate was calculated during the shaded epoch of the recording following threshold crossing at 0 mV. The subthreshold membrane voltage relationship obtained before blue light stimulation is highlighted in orange and superimposed onto the voltage response recorded after blue light stimulation for comparison. In this case the subthreshold response does not change following light stimulation. ***B***, Plots illustrating the change in average AP rate in the FS-IN_*na*_ population (black symbols) and the FS-IN_*a*_ population (green symbols). The time course of these changes in average AP rate in the FS-IN_*na*_ population can be described using a Boltzmann function and the shaded area indicates the 95% confidence limits for these fits. ***C*** & ***D***, same conventions as ***A*** & ***B*** but the data was obtained from the FS-IN_*na*_ population in the presence of H1/H2 antagonists. Note in ***C*** how the subthreshold I-V relationship of FS-INs does not alter following optogenetic stimulation of TMN_HDC_ axons and how in ***D*** there is no increase in AP rate following stimulation.

### Age differences associated with TMN_HDC_ modulation of the PFC

We next explored the age-dependence of TMN_HDC_ modulation of the PFC with a focus on changes in the input conductance of PyrNs. First, we examined if there were any age or sex related differences in the resting input conductance of PyrNs or FS-INs. As shown **Figure 5A**, the resting input conductance of PyrNs was similar across the adult lifespan and this parameter was not influenced by an animal’s sex. A linear regression analysis of this data (male & female) resulted in a negative slope and a Pearsons’s correlation coefficient or r^2^ of - 0.03, indicating that variability in the resting input conductance was not significantly influenced by the age of the animal. Across the entire lifespan, the resting input conductance of the PyrNs recorded from male mice was 4.44 ± 0.36 nS (n=55) compared to 5.44 ± 0.47 nS (n=46) for female mice.

**Figure 5.**
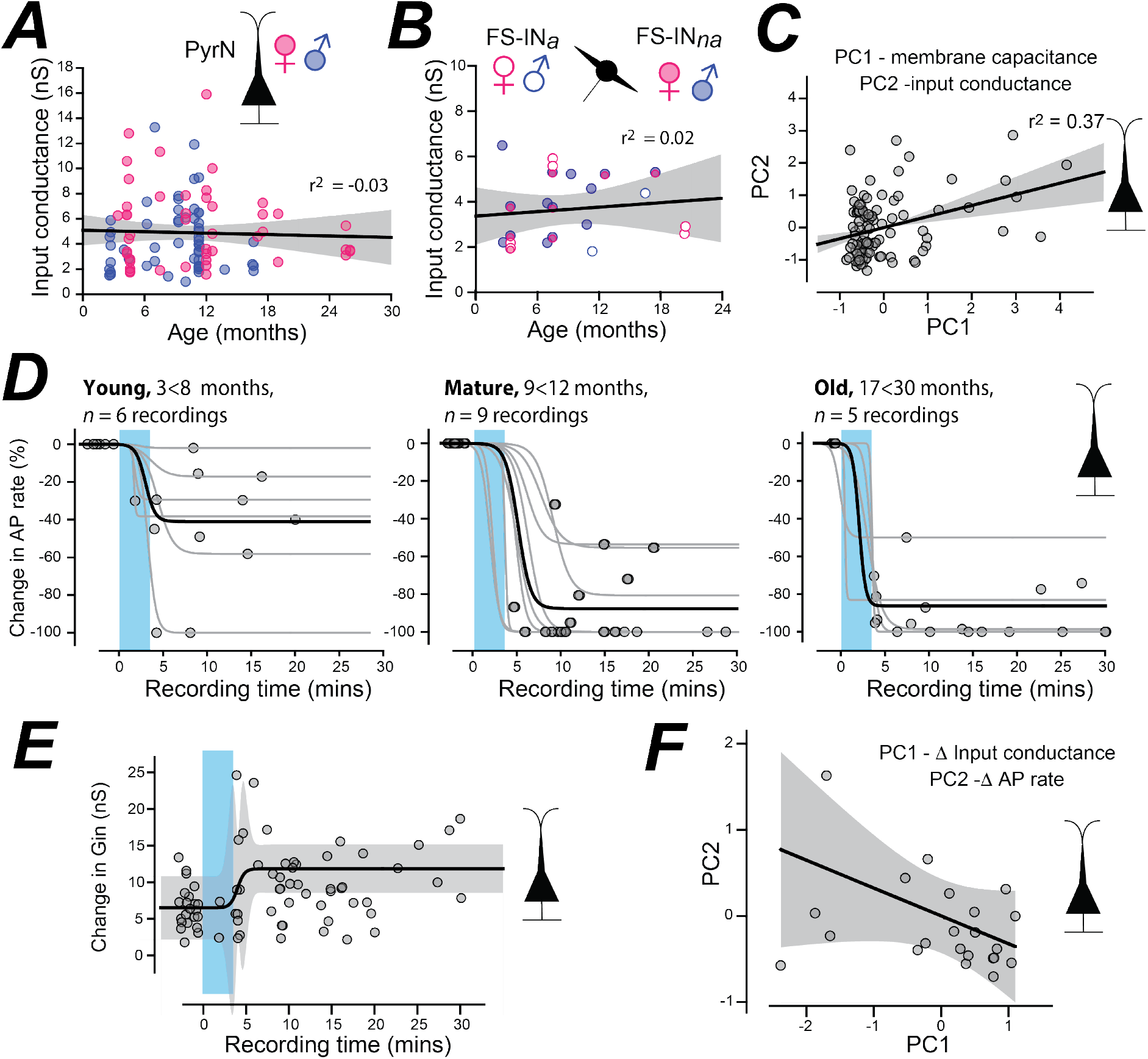
Age- and sex-related differences in PFC excitability changes. ***A***, Scatter plot of the resting input conductance taken from all PyrNs recorded from mice aged 3 months to 27 months. This data is separated into male (blue circles) and female (pink circles). The solid line is the result of linear regression analysis to the combined male and female data demonstrating a lack of change in input conductance across the adult lifespan. The grey shaded area shows the 95% confidence level for this fit and the r^2^ value of −0.03 indicates how little of the variability in resting excitability can be explained by age. ***B***, Scatter plot like ***A*** containing data from FS-IN cells. Measurements from FS-IN_a_ and FS-IN_na_ cells have been separated into male and female but once again the linear region indicates that the resting excitability of FS-INs does not vary across the adult lifespan. ***C***, Normalised data comparing the relationship between the membrane capacitance (PC1) and the input conductance (PC2) for all PyrNs recorded across all ages. The linear regression analysis demonstrates that there is some relationship between these two parameters. ***D***, A series of time course plots taken from PyrNs at three age ranges. These plots illustrate the change in AP rate before and after optogenetic stimulation of TMN_HDC_ axons. Each point represents the change in the average AP rate calculated during current injection experiments before and after the optogenetic stimulation (blue shaded area). These experiments are identical to those illustrated in Figure 3. Once again, optogenetic stimulation consisted of 4 minutes of 5 Hz blue light bursts of 1 ms duration delivered every 0.4 seconds. All AP rate changes are relative to the initial value obtained immediately prior to stimulation. The dark line in each plot is the average of all sigmoidal functions fitted to each experiment. ***E,*** Time course of the change in input conductance recorded for PyrNs across all ages. The sigmoidal fit describes the consistent increase in the input conductance that is observed following TMN_HDC_ stimulation. The shaded area illustrates the 95% confidence levels for this fit. ***F,*** Normalised data comparing the relationship between the change in the input conductance (PC1) and the change in AP rate (PC2) for all PyrNs recorded across all ages. The linear regression analysis demonstrates that there is some relationship between these two parameters such that as input conductance increases the AP rate reduces. ***G***, Scatter plot of the resting input conductance for male (blue) and female (pink) PyrNs in the PFC before and after optogenetic stimulation of the TMN_HDC_ axons. The straight lines illustrate the results of linear regression analysis demonstrating how the change in input conductance is greater for the female data compared to the male data.

Much of the variability observed for the resting input conductance in the PyrN population (4.90 ± 0.36 nS, *n* = 101) can be explained by variability in cell size as determined from the membrane capacitance (39.1 ± 3.6 pF). A least square regression analysis (**Figure 5B**) of the normalised data demonstrates a slope of +0.34 with an r^2^ value of 0.37 indicating that nearly 40% of the variability in the input conductance (PC2) can be explained by differences in cell size (PC1). The resting input conductance of FS-INs (n = 23 recordings) was also not related to the age of the mouse with no correlation indicated following linear regression analysis (r^2^ = 0.02). Although the number of recordings in each group was too small to attribute any statistical significance to these changes, the resting input conductance of FS-IN_na_ and FS-IN_a_ populations were similar in magnitude and were also not influenced by sex or age (**Figure 5C**).

We did observe changes in the response of PyrNs to the release of GABA/histamine from TMN_HDC_ axons across the adult lifespan (**Figure 5D**). Differences between the three age groups were observed at p < 0.02 using Kruskal Wallis ANOVA, and then further Mann Whitney tests revealed a significant difference between the ‘young’ and ‘mature’ age groups with n_young_= 6, n_mature_= 9, p < 0.008. Differences between the change in excitability observed in the three age groups was present at p < 0.02 using Kruskal Wallis ANOVA. Mann Whitney tests revealed a significant difference between the ‘young’ (from 3 months and up to 8 months old) and ‘mature’ (from 8 months and up to 11 months old) age groups with n_young_= 6, n_mature_= 9, p < 0.008. The time course for the rapid reduction in PyrN excitability was described by a simple Boltzmann function that indicated that the reduction in excitability took place 2 to 5 minutes following the start of blue light stimulation at a rate of over −50% per minute. More importantly, these fits demonstrated how the ability of TMN_HDC_ axons to reduce PyrN excitability was greater in older animals (**Figure 5D**).

Consistent with the modulation of a tonic GABA_A_ receptor mediated conductance following GABA release from TMN_HDC_ axons, we observed a clear enhancement of the input conductance in PyrN recordings that followed the same time course as the reduction in AP firing (**Figure 5E**). The fit to the data recorded from PyrNs across all ages showed an increase in the input conductance from 6.52 ± 2.16 nS to 11.85 ± 1.65 nS that occurred 4 minutes after light stimulation began. The increase in the input conductance did correlate with the magnitude of the reduction in AP firing across the PyrN population, as shown by the linear regression analysis of the normalised data with an r^2^ value for this fit of −0.323 (**Figure 5F**). Therefore, the age-related changes we observe in TMN_HDC_ modulation of AP firing can be explained by enhanced modulation of the input conductance in later life.

## Discussion

This study has demonstrated how histamine/GABA released from TMN_HDC_ axons alters information transfer within the mouse PFC in a cell- and age-specific manner. We recorded from three distinct classes of PFC neuron in prelimbic (PL), anterior cingulate (AC) and infralimbic (IL) regions of the PFC: layer 2/3 pyramidal neurons (PyrN), non-accommodating fast-spiking interneurons (FS-IN_na_) and accommodating fast-spiking interneurons (FS-IN_a_). Each cell-type responded differently to optogenetic stimulation of TMN_HDC_ axons. The FS-IN_a_ population was unaltered while the FS-IN_na_ population (parvalbumin-like cells) was excited and the PyrN population was robustly inhibited following TMN_HDC_ activation (see conceptual summary **Figure 6**). Pharmacological experiments confirmed that histamine release was responsible for FS-IN_na_ excitation, whereas the release of GABA from THN_HDC_ axons was responsible for the inhibition of PyrNs. By making recordings from mice over an extended period of their adult lives (3-27 months postnatal), we found that the shunting inhibition produced by this GABA modulation was more pronounced in the adult and was sustained into older age.

**Figure 6.**
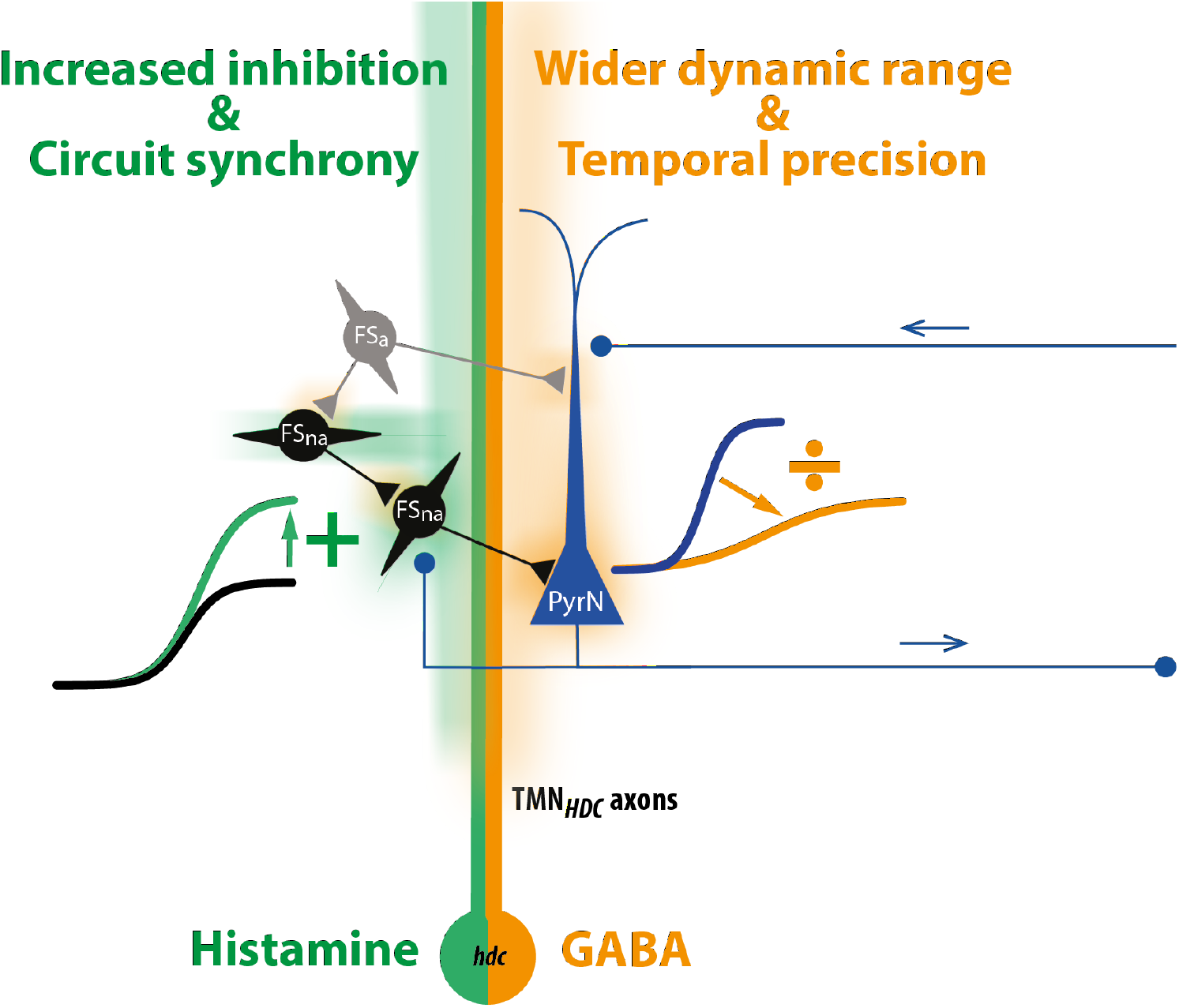
Summary of salient features associated with TMN modulation of the PFC. Illustration that highlights the actions of the histamine (green) and GABA (orange) released from TMN_HDC_ axons. The GABA released from TMN_HDC_ axons will lead to enhanced precision and greater dynamic range within the PFC due to the divisive gain change produced within the PyrNs. In parallel, the histamine coreleased from TMN_HDC_ axons will increase synchrony within the PFC network due to the additive gain change associated with the FS-IN_na_ population.

Histamine supports many aspects of wakefulness, so although the net effect of the histamine system in the brain is excitatory at the behavioural level (Haas and Panula, 2003; Scammell et al., 2019; Yoshikawa et al., 2021), in the case of the neocortical network we propose that the histamine-GABA system influences the circuits requirement for AP precision. On the one hand, we find that histamine stimulates a subset of fast-spiking GABAergic interneurons that will increase phasic inhibition onto PyrNs to ultimately enhance precision of PyrNs and expand their dynamic range to enhance synchrony in the PFC network (Krimer et al., 2005; Feldmeyer et al., 2018). On the other hand, the raised ambient GABA levels produced following TMN_HDC_ axon GABA release will speed up the membrane time constant of PyrNs by enhancing the resting input conductance following extra synaptic GABA_A_ receptor activation and, therefore, will also enhance coincidence detection, such that more closely timed EPSP inputs will be required to elicit APs (Brickley and Mody, 2012; Wlodarczyk et al., 2013; Sylantyev et al., 2020). By limiting the window of coincidence detection, tonic inhibition can be seen to promote conditions that enhance cognition. Moreover, we propose that the additive gain changes associated with the FS-IN_na_ population of the PFC will further promote synchrony of the network during wakefulness.

Previously, gain changes within neocortical circuits have mostly been studied in relation to the action of neuromodulators such as 5HT and acetylcholine on local interneurons (Ferguson and Cardin, 2020). However, this is different to the mechanism we currently propose, whereby release of GABA from TMN_HDC_ axons is directly responsible for the generation of a tonic shunting inhibition (**Figure 6**). Within minutes of TMN_HDC_ opto-activation, the PyrN population experienced a dramatic reduction in the slope of the input-output relationship and a reduction in the maximum AP firing rate. As expected, this divisive gain change was associated with a significant 20% increase in the input conductance, with no change in the RMP, a characteristic of a tonic shunting inhibition mediated by extrasynaptic GABA_A_ receptor activation (Mitchell and Silver, 2003). Therefore, GABA diffusion through the extracellular space is sufficiently unhindered to enable rapid coupling between the GABA released from TMN_HDC_ axons and alterations in the gain of the surrounding PyrNs. This is similar to the mode of action reported in other cortical and striatal regions following TMN_HDC_ activation (Yu et al., 2015), and consistent with observations following the knockout of high-affinity δ subunit-containing GABA_A_ receptors (Abdurakhmanova et al., 2020).

We have also investigated the consequence of TMN_HDC_ activation for local interneuron excitability and observed an increase in the gain of these neurons, but this gain change was predominantly additive in nature, and the time course of the response was much slower than that observed for PyrNs. In contrast to the effects of TMN_HDC_ activation on PyrNs, the additive gain change observed for the FS-IN_na_ population was due to histamine’s actions on H1/H2 receptors. The delayed increase in FS-IN_na_ excitability is consistent with the reduction of an outward potassium conductance that has been reported in hippocampal interneurons following H2 receptor activation and the subsequent coupling to adenyl cyclase pathways (Atzori et al., 2000). However, the divisive gain change that results from the direct actions of TMN_HDC_ axon-released GABA on extrasynaptic GABA_A_ receptors clearly dominates in the PFC, as this still occurred in the presence of H1/H2 receptor blockers when the excitability of surrounding interneurons was not altered.

Human brain imaging studies have reported lower levels of variability in the resting state BOLD signal in the PFC of individuals in later life (Nomi et al., 2017), and this reduction in signal variability has been associated with overall decreased GABA levels in the cortex (Porges et al., 2021). In older humans, positive allosteric modulators of γ2 subunit-containing GABA_A_ receptors, for example with lorazepam, may protect against cognitive decline (Lalwani et al., 2021). Tonic (extra-synaptic) inhibition can come from GABA-activation of both γ2 and δ subunit-containing receptors, but only γ2 subunit-containing GABA_A_ receptor mediated responses can be allosterically enhanced by benzodiazepines. Similarly, for individuals that maintain cognitive performance with aging, the increases in TMN_HDC_ GABA modulation of PyrNs during ageing may be an adaptive mechanism to buttress cognition.

Finally, we should note that our present and previous results on GABA-histamine corelease do differ from results reported in another study (Venner et al., 2019). In general, most findings, whether pharmacological or genetic, support the role of histamine in promoting aspects of wakefulness, although, as usually found, permanent gene or cell knockouts/lesions tend to give different results from reversible pharmacology or knockdowns: for example, *hdc* gene knockouts, and *hdc* cell lesions induce sleep-wake fragmentation (insomnia), but do not affect overall levels of sleep-wake (Parmentier et al., 2002; Yu et al., 2019); on the other hand, histamine H1 receptor antagonists, chemogenetic or opto-inhibition inhibition of histamine neurons induces NREM-like sleep (Fujita et al., 2017; Yu et al., 2019; Naganuma et al., 2021; Yoshikawa et al., 2021). We have previously explored how levels of arousal are altered following chemogenetic manipulation of this TMN_HDC_ projection (Yu et al., 2015), and found that activation of the TMN_HDC_ projection increased motor activity, and that genetic knockdown of *vgat* from this pathway sustains wakefulness. We do not know why our work differs from the findings of Venner et al. 2019, who found, that chemogenetic manipulation of hdc cells did not affect sleep-wake behaviour and found no role for GABA in wakefulness, since after knockout of *gad* and *vgat* genes from *hdc* cells there was no behavioural phenotype (Venner et al., 2019; Arrigoni and Fuller, 2021).

In the adult, the TMN is the only location where the *hdc* gene is expressed neuronally (Watanabe et al., 1983; Panula et al., 1984; Bayliss et al., 1990). The *hdc* gene therefore allows selective genetic manipulations of histamine neurons. At least three Cre recombinase mouse lines based on the *hdc* gene are available (Yanovsky et al., 2012; Zecharia et al., 2012; Fujita et al., 2017). The line we developed is a *knock-in* of the Cre gene into the *hdc* gene and in the adult has expression only in histamine cells and ventricular ependyma in the TMN (Zecharia et al., 2012); the other lines are based on BAC *Hdc-Cre* transgenes randomly integrated into the genome (Yanovsky et al., 2012; Fujita et al., 2017), and do not express Cre recombinase in all *hdc* neurons.

The Cre line used by Venner et al does not target all *hdc*-expressing neurons. For our mechanism of GABA released, we propose the mechanism is via VGAT (Yu et al., 2015), but in any case, GABA can also be transported into synaptic vesicles by the vesicular monoamine transporter VMAT (Tritsch et al., 2012), whose gene is strongly expressed in histaminergic cells (Mickelsen et al., 2020). Venner et 2019 did not provide electrophysiological evidence to support their contention that GABA was not co-released from TMN_HDC_ axons in the neocortex, although they found using slice electrophysiology that histamine axons projecting to the preoptic hypothalamus did not release GABA. The best explanation for the disparities between studies are that the various subtypes of histamine neuron identified by RNAseq studies differ in their ability to release GABA (Mickelsen et al., 2020).

In summary, we present evidence that histamine released from TMN_HDC_ axons is responsible for an additive gain change mechanism within specific interneuron populations of the adult PFC. We speculate in **Figure 6**, that these changes in FS-IN_na_ excitability will lead to increased synchronisation of cortical circuitry of the type (high frequency) associated with the awake brain (Yu et al., 2015). In contrast, the GABA released from these same TMN_HDC_ axons leads to a divisive gain change that will broaden the dynamic range of PyrNs (**see Figure 6**) a feature that will lead to greater computational flexibility within the PFC circuitry. We have also shown that the shunting inhibition associated with TMN_HDC_ axons increases with age and, it is intriguing to speculate that the enhancement of this feature of TMN_HDC_ modulation could be a compensation for age-related declines in global GABA levels observed within the PFC (Porges et al., 2021). Enhanced GABA modulation from TMN_HDC_ axons could help protect against the reduced cognitive flexibility that is a feature of the ageing process while the histamine that is released will support increased arousal due to modulation of local interneurons.

## Acknowledgments

Funded by the Wellcome Trust (107841/Z/15/Z, W.W.), the UK Dementia Research Institute (UK DRI-5004, W.W.) and the BBSRC (P.C and S.G.B, BB/N008871/1)

